# Pancreatic Deletion of Mitogen-inducible Gene 6 Promotes Beta Cell Proliferation Following Destruction

**DOI:** 10.1101/2023.01.30.526325

**Authors:** Kimberley El, Brandon M. Bauer, Yi-Chun Chen, Jae-Wook Jeong, Patrick Fueger

**Affiliations:** Department of Cellular & Integrative Physiology, Indiana University School of Medicine, Indianapolis, IN 46202; Department of Molecular and Cellular Endocrinology, Beckman Research Institute of the City of Hope, Duarte, CA 91010; Department of Obstetrics, Gynecology and Women’s Health, University of Missouri School of Medicine, Columbia, MO 65211

## Abstract

Type 1 Diabetes (T1D) is caused by autoimmune-mediated beta cell destruction. Following beta cell injury, the pancreas attempts to launch a cellular repair and regenerative program, yet it fails to completely restore functional beta cell mass. One component of this regenerative program is epidermal growth factor receptor (EGFR) signaling. However, upon irreparable beta cell damage, EGFR signaling is dampened, disrupting attempts to restore functional beta cell mass and maintain normoglycemia. We previously demonstrated that the negative feedback inhibitor of EGFR, Mitogen-inducible gene 6 (Mig6), is induced by the pro-inflammatory cytokines central to the autoimmune-mediated beta cell destruction. We also established that pro-inflammatory cytokines suppress EGFR activation, and siRNA-mediated suppression of Mig6 restores EGFR signaling. Thus, we hypothesized that pro-inflammatory cytokines induce nitric oxide production and that in turn induced Mig6, disrupting EGFR repair mechanisms. We determined that NO induces Mig6, attenuating EGFR signaling, and NO synthase inhibition blocks the cytokine-mediated induction of Mig6, thereby restoring cytokine-impaired EGFR signaling. To that end, we treated mice lacking pancreatic Mig6 and control mice with a streptozotocin (STZ) to induce beta cell death and diabetes in a way that mimics the onset and progression of T1D. Whereas STZ-treated control mice became hyperglycemic and had reduced beta cell mass, STZ-treated Mig6 pancreas-specific knock out (PKO) mice remained euglycemic and glucose tolerant due to preserved beta cell mass. The restoration of beta cell mass in PKO mice was accompanied by enhanced beta cell proliferation. Thus, our work suggests that Mig6 is a promising target to preserve beta cell mass before overt T1D.

## INTRODUCTION

Type 1 diabetes (T1D) is a progressive disease characterized by autoimmune-mediated destruction of the pancreatic insulin-secreting beta cells. Multiple human and animal studies support the concept that progression to overt diabetes is accompanied by both beta cell dysfunction and destruction [1, 2]. Beta cell destruction in T1D can occur through a series of immunological attacks and recoveries, termed “relapsing-remitting” [3]: a process which gradually reduces functional beta cell mass until symptoms of hyperglycemia are noticed and diabetes is clinically diagnosed. During the progression to diabetes, immune cells such as macrophages are recruited and promote inflammation by releasing pro-inflammatory cytokines into the islet [4], which activates stress activated kinases, as well as NF-κB signaling and iNOS expression [5, 6]. Such signaling events lead to increased endoplasmic reticulum (ER) stress, increased apoptosis, and decreased beta cell function and regeneration, all of which culminates in the loss of functional beta cell mass and diabetes.

The relapsing-remitting features of diabetes extend past clinical diagnosis to the ‘honeymoon phase’, where residual beta cell mass provides endogenous insulin to support exogenously delivered insulin by the patient. This honeymoon phase provides an opportune time for therapeutic intervention; in fact, numerous clinical trials for delivering immunomodulatory agents in this window have been reported. These therapeutic strategies targeting the immune response but neglecting intrinsic cellular pathologies within the islet have resulted in immunological approaches that are largely ineffective in the long term. In contrast, cellular therapies can promote beta cell proliferation, survival, and recovery, essentially restoring beta cell mass [7]. Nevertheless, translating pre-clinical successes to patients with T1D has been challenging. For example, administration of epidermal growth receptor (EGF) and gastrin in combination increases beta cell mass and reverses hyperglycemia in NOD mice [8], but this approach has yet to translate into a viable treatment for humans.

The EGF receptor (EGFR) cascade has proven to be crucial in the formation and regulation of pancreatic beta cell mass as it modulates cell growth, proliferation, survival, and differentiation [9]. EGF ligand is routinely combined with Notch inhibitors to convert stem cell-derived pancreatic progenitor cells into pancreatic endocrine progenitor cells [10–12]. Mice lacking EGFR acquire diabetes within two weeks of birth and have impaired islet development [13], and mice expressing constitutively active EGFR have increased beta cell mass [14]. Further, heparin-binding EGF-like growth factor (HB-EGF), an EGFR ligand, promotes rodent beta cell proliferation *in vivo* in response to nutrient excess and human beta cell proliferation *in vitro [15]*. EGFR signaling activates downstream effectors such as Akt, which promotes cell survival, and ERK, which increases cell proliferation [16]. Additionally, in some beta cell regeneration studies expression of EGF ligands increases [17, 18], suggesting that this pathway is involved in an intrinsic beta cell regeneration and repair program.

However, endogenous feedback mechanisms restrain EGFR activation, thereby preventing the full regenerative actions of EGFR signaling. One such mechanism is Mig6, a cellular response protein and feedback inhibitor of EGFR that has been characterized as a molecular “brake” for beta cell proliferation [16, 19] and survival [1, 16]. Activation of EGFR signaling induces Mig6 in a classic feedback mechanism, suppressing kinase activity and initiating receptor endocytosis and degradation [20]. Mig6 haploinsufficient mice are protected against chemically-induced diabetes [1], suggesting that Mig6 antagonizes beta cell mass. In addition, increasing Mig6 expression with a recombinant adenovirus compromises beta cell integrity and islet function [1]. Thus, Mig6 expression levels can modulate functional beta cell mass.

Not only is Mig6 induced during EGFR activation, but it is also activated by other factors, such as the pro-inflammatory cytokines from the T1D milieu [1]. Pro-inflammatory cytokines are well established mediators of beta cell damage [21, 22], and this concept has been verified *in vitro* [23]. Previous work has determined that of all the cytokines released during the immune response, only TNF-α, IL-1β, and IFN-γ are required to produce the full inflammatory response [24]. TNF-α and IL-1β stimulate their respective receptors and initiate a downstream signaling cascade that results in transcription of inducible NO synthase (iNOS) and consequent NO production [25–27]. NO is essential for beta cell dysfunction and apoptosis [6]. Besides activation of the NF- κB pathway and NO production, we have demonstrated that cytokines reduce the EGFR signaling, which can be rescued by Mig6 suppression [1]. Here, we demonstrate that NO is necessary but insufficient to induce Mig6 during exposure to pro-inflammatory cytokines. Given that Mig6 antagonizes the actions of EGFR signaling, we hypothesized that reducing Mig6 expression in the pancreas would provide protection in a beta cell destruction model by promoting beta cell proliferation.

## MATERIALS AND METHODS

### Cell experiments

INS-1-derived 832/13 rat insulinoma cells were cultured as previously described [28]. A starvation medium (RPMI 1640 containing 2.5 mmol/l glucose and 0.1% BSA) was used for EGF stimulation experiments. For cytokine plus EGF stimulation experiments, 832/13 cells were pretreated with a pro-inflammatory rat cytokine ‘cocktail’ (1000 U/ml TNFa, 50 U/ml IL-1β, and 1000 U/ml IFNγ; Prospec, East Brunswick, NJ) for 16 hours, starved for 2 hours and treated with 10 ng/ml rat recombinant EGF (R&D Systems, Minneapolis, MN, USA) for 5 min. For nitic oxide experiments, 832/13 cells were treated with L-NMMA (MilliporeSigma, St Louis, MO, USA) for 16 h, or DPTA/NO (Cayman Chemical, Ann Arbor, MI, USA) for 4 h.

### Immunoblot analysis

Immunoblot analysis was performed as previously described [16]. Phosphorylated EGFR^Y1068^ (Cell Signaling, #3777, 1:1000 with Nacalai USA Signal Enhancer HIKARI) levels were normalized to total protein levels (Sigma-Aldrich, #E3138, 1:1000 with Nacalai USA Signal Enhancer HIKARI), and total (i.e. nonphosphorylated) protein levels were normalized to tubulin protein levels (Sigma-Aldrich, #T6557, 1:2500 in TBS with 2% polyvinylpyrrolidone).

### Quantitative RT-PCR analysis

RNA from 832/13 cells, and mouse, rat, and human islets was isolated using RNeasy Mini or Micro kits (Qiagen, Valencia, CA, USA). Reverse transcription was performed using a High Capacity cDNA Reverse Transcription kit (Applied Biosystems, Foster City, CA, USA). Threshold cycle ΔCT methodology was used to calculate the relative quantities of *Errfi1*, *Nos2*, and *Gapdh* (TaqMan assays, Applied Biosystems) mRNA [29]. Primer sequences were described previously [30]. PCR reactions were performed in triplicate for each sample from at least three independent experiments and normalized to *Gapdh* mRNA levels.

### Animals and treatments

All animals were maintained and used in accordance with protocols approved by the Institutional Animal Care and Use Committees at Indiana University School of Medicine or the City of Hope, following the *Guide for the Care and Use of Laboratory Animals*, Eighth edition (2011). Mice were maintained in a standard 12-hour light-dark cycle and provided unrestricted access to water and a standard rodent chow. C57Bl/6J mice, with the Pdx-1 promoter controlling element expression of Cre recombinase in the pancreas (Jackson Laboratory #014647), were bred with *Mig6^flox/flox^* or *Mig6*^flox/+^ mice on a C57Bl/6J background [31]. DNA was extracted from a tail biopsy and genotypes were determined using the Extract-N-Amp Tissue PCR kit (Sigma-Aldrich, St Louis, MO, USA). The following primer sequences (Integrated DNA Technologies) were used to detect Pdx1-Cre and *Mig6^flox^*: Pdx-forward = 5’-CTGGACTACATCTTGAGTTGC, Pdx-reverse = 5’-GGTGTACGGTCAGTAAATTTG, *Mig6*^flox^-forward = 5’-GGTCAGGGCTGTGCAGTCCGTAGA, *Mig6*^Neo^-reverse = 5’-CGATACCCCACCGAGACC, and *Mig6*^flox^-reverse = 5’-CTTCCCAAATCTAACACCCGACAC.

Eight to ten-week old male mice of the genotypes Mig6^flox/flox^/Cre^+^ (Mig6 pancreatic knock out, PKO), and Mig6^flox/flox^/Cre^-^ (controls) were intraperitoneally injected with streptozotocin (STZ, 35mg/kg body weight; Sigma-Aldrich, St Louis, MO, USA) for 5 consecutive days as performed previously [1]. A group of control animals was injected in the same manner with vehicle (saline).

### Immunohistochemical and immunofluorescence staining

Immunostaining of pancreatic sections was performed as previously described [1]. Antibodies are listed in Supplementary Data Table 1. Beta cell cross-sectional area relative to total islet area was used as a surrogate for quantifying beta cell mass.

### Insulin secretion *in vivo*

Hyperglycemic clamps were used to quantify insulin secretion *in vivo*. Surgical procedures have been described in detail previously [32, 33]. Briefly, mice were anesthetized with isoflurane, and the carotid artery and jugular vein were catheterized. Free catheter ends were tunneled under the skin to the back of the neck and externalized with a MASA^TM^ (Mouse Antenna for Sampling Access), which permits arterial sampling from an indwelling catheter in conscious mice without handling stress. After surgery, mice were individually housed and recovered for up to 5 days to allow for body weight to return to within ~10% of presurgical body weight.

On the day of study, mice were fasted for 5 h and connected to a swivel. Saline-washed erythrocytes were infused (5–6 μl/min) during the experimental period to prevent a >5% fall in hematocrit. Blood samples were collected from the arterial catheter in tubes containing EDTA and centrifuged, and plasma was stored at −20°C until analyzed. At t = 0 min, a variable glucose infusion rate (GIR) was initiated to increase and maintain blood glucose at ~15.0 mmol/l, with a 100 mg/kg/min rate used for 2 min to provoke first-phase insulin secretion. Blood glucose (5 μl) was measured at t = –15, –5, 5, 10, 15, and 20 min and then every 10 min until t = 120 min with an Alpha Trak glucometer (Abbott Laboratories, Abbott Park, IL, USA). Larger blood samples (100 μl) to measure plasma insulin and C-peptide were taken at various timepoints.

### Glucose tolerance testing

For glucose tolerance tests (GTT), 1.5 g/kg body weight D-glucose (Sigma-Aldrich) was intraperitoneally injected into 6-h-fasted saline-control or STZ-treated mice. Blood was sampled from a tail vein at the indicated time points, and blood glucose was measured using an AlphaTrak glucometer.

### Statistical analysis

All data are presented as means ± SEM. Protein and mRNA data were normalized to control conditions and presented as relative expression. Adjusted area-under-the-curve (AUC) was calculated using the trapezoid method. Student’s *t*-test (unpaired, two-tailed unless otherwise stated) or ANOVA (with Bonferroni post hoc tests) were performed using GraphPad Prism software (La Jolla, CA, USA) to detect statistical differences. *p < 0.05* was considered statistically significant.

## RESULTS

### NO is detrimental to EGFR signaling

Previous studies in our lab have reported that cytokines dampen EGFR phosphorylation and that Mig6 suppression by siRNA rescues this inhibition in 832/13 cells [1]. Given our findings on the relationship between NO and Mig6, we investigated the direct role of NO on EGFR signaling. As an extension of our previous work [1], we used DPTA as an NO donor and discovered that NO alone, like cytokines, dampens EGFR phosphorylation (**Figure 1A-C**).

**Figure 1.**
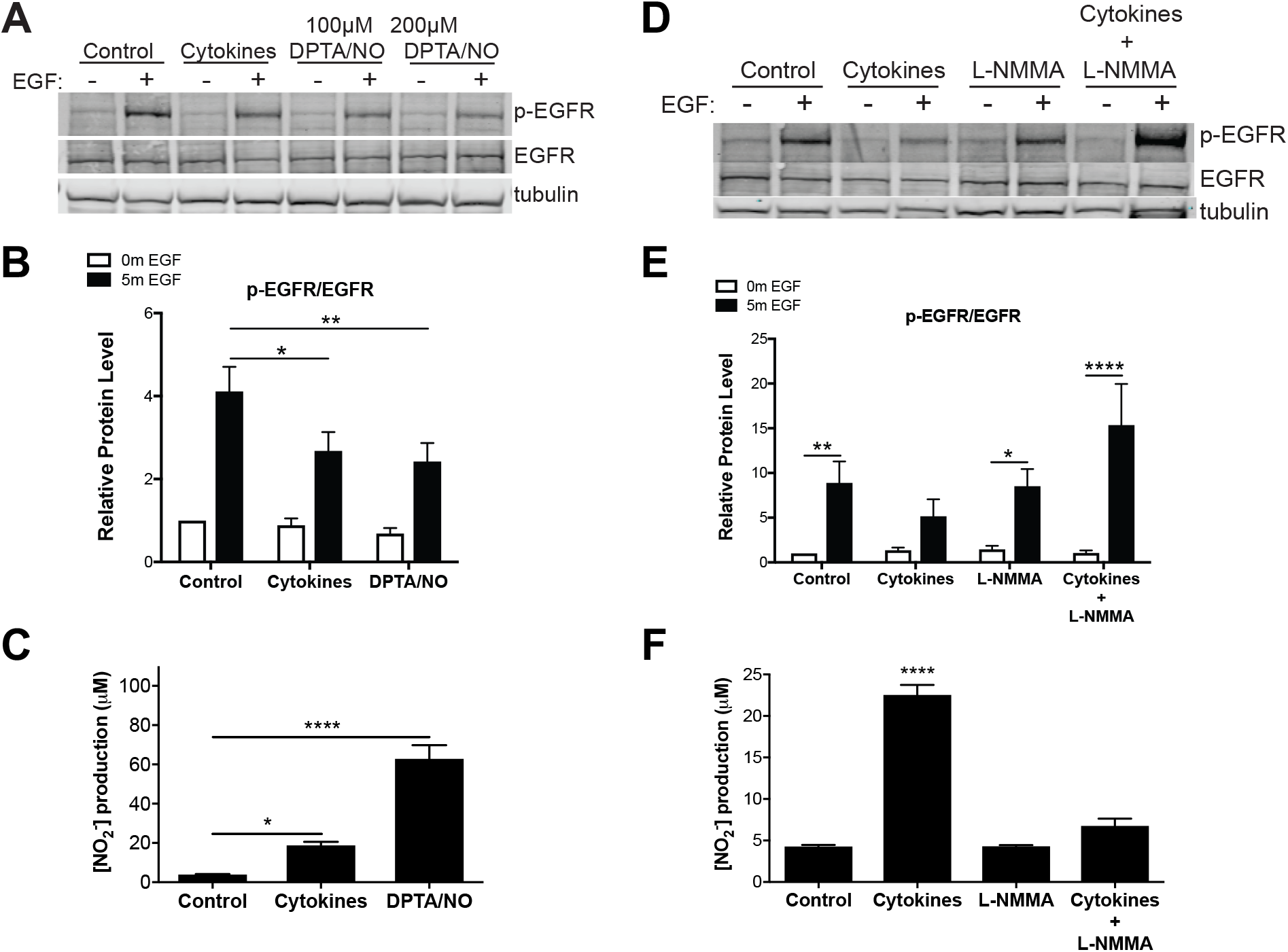
Cytokines and nitric oxide attenuate EGFR signaling. **(A)** Representative image of western blot in which 832/13 cells were treated with cytokines (TNF*α*, IL-1*β*, IFN-*σ*; 8h) or nitric oxide donor, DPTA/NO (4h), starved for 2h in 2.5mM glucose and stimulated with 10 ng/ml rrEGF for 5 min. Cell lysates were collected for western blot analysis to determine p-EGFR, EGFR, and tubulin levels. **(B)** Quantified p-EGFR/EGFR reported as protein levels relative to non-cytokine, non-EGF treated groups and normalized to tubulin. *n = 4-5;* *, *p<0.05 vs control EGF-stimulated;* **, *p<0.01 vs control EGF-stimulated, 2-way ANOVA*. **(D)** Concentration of NO_2_^-^ in the media was measured using Griess assay. *n=4;* *, *p<0.05;* ***, *p<0.001 vs. untreated control, 1-way ANOVA*. **(D)** Representative image of western blot where 832/13 cells were treated with cytokines (TNF*α*, IL-1*β*, IFN-*γ*; 16h) and/or iNOS inhibitor, L-NMMA (16h), starved for 2h in 2.5mM glucose and stimulated with 10 ng/ml rrEGF for 5 min. Cell lysates were collected for western blot analysis to determine p-EGFR, EGFR, and tubulin levels. **(E)** Quantified p-EGFR/EGFR shown as protein levels relative to non-treated groups and normalized to tubulin. *n = 7;* *, *p<0.05;* **, *p<0.01;* ***, *p<0.001;* ****, *p<0.0001 vs non-EGF-stimulated, 2-way ANOVA*. **(F)** Concentration of NO_2_^-^ in the media was measured using Griess assay. *n=3;* ****, *p<0.0001 vs. untreated control, 1-way ANOVA*.

To further assess the role of NO in EGFR signaling, we treated 832/13 cells with the iNOS inhibitor, L-NMMA, and observed that cytokines dampen EGFR signaling as expected, but that a combinatory treatment of cytokines and L-NMMA caused an overwhelming restoration of EGFR signaling (**Figure 1D-F**). Taken together, the NO donor and iNOS inhibitor data indicate that NO is detrimental to EGFR signaling.

### Cytokine-induced Mig6 expression requires NO

It is widely accepted that pro-inflammatory cytokines, the mediators of immune-cell damage to beta cells, are involved in the progression to T1D [34]. Pro-inflammatory cytokines induce Mig6 expression in both human islets and 832/13 cells [1]. Here, we verify that cytokines increase Mig6 expression in the 832/13 cell line (**Figure 2A**), demonstrating that cytokine-induced Mig6 expression requires cytokine-mediated NO. This induction of Mig6 with cytokines was prevented by co-treatment with L-NMMA. Interestingly, NO alone was not sufficient to increase Mig6 expression in either a dose- or time-dependent manner (**Figure 2B,C**), suggesting NO functions in a permissive manner to induce Mig6 with activated cytokine signaling.

**Figure 2.**
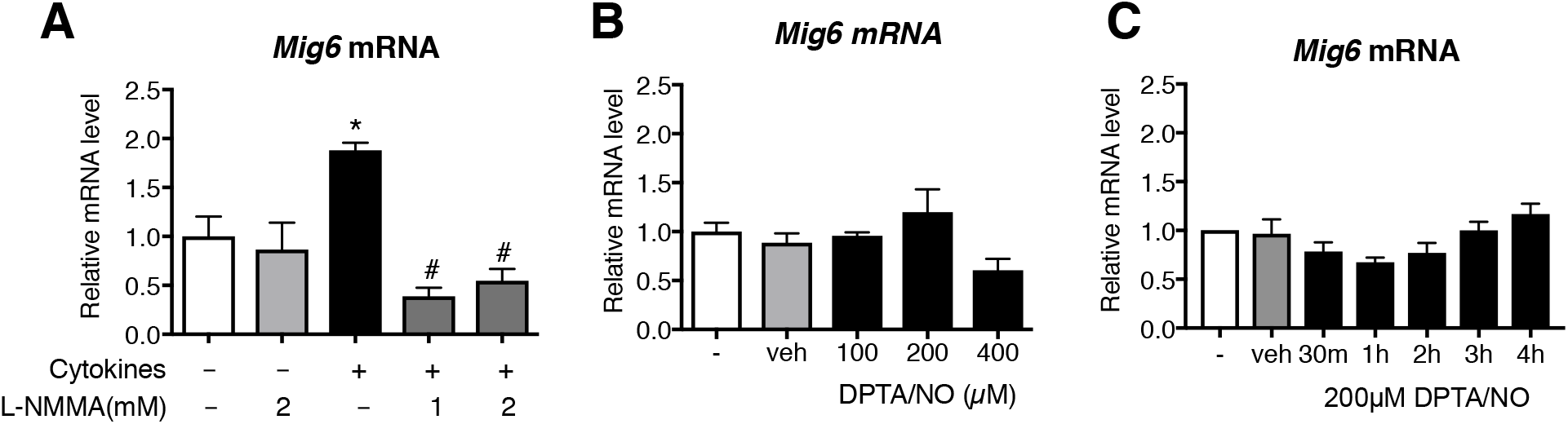
*Mig6* expression requires cytokine-induced NO, but NO alone is insufficient. **(A)** 832/13 cells were treated with cytokines (TNF*α*, IL-1*β*, IFN-*γ*) and/or the iNOS inhibitor, L-NMMA for 24 h or **(B,C)**DPTA/NO at 100-400μM at times ranging from 30min to 4h. *Mig6* and *Gapdh* mRNA levels were determined by qRT-PCR. *n = 3-6;* *, *p<0.05 vs. non-treated control, Student’s t-test;* ^#^, *p<0.05 vs. cytokine-treated, 1-way ANOVA*.

### Mig6 PKO mice have normal beta cell mass and function

As previously reported, whole-body Mig6 heterozygous knock-out mice were protected from streptozotocin (STZ)-induced glucose intolerance, with no change in insulin sensitivity [1]. Yet, this mouse model was not pancreas-specific, and the corrections in glucose tolerance waned at 20-days post STZ injection. To directly assess Mig6’s actions in the pancreas, we developed PKO mice by breeding a *Pdx-Cre* mouse model with *Mig6^flox/flox^* mice, developed as previously described [31]. Compared to control mice, PKO mice exhibited a 50% knock down of Mig6 protein and 60% knock down of Mig6 mRNA in the pancreatic islet (**Figure 3A-C**). Control and Mig6 PKO mice have no difference in beta cell mass (**Figure 3D**), quantified from immunohistochemical (IHC) staining of insulin as insulin-positive area relative to pancreas area. We also observed no major differences in islet morphology (**Figure 3E**). Using the gold-standard hyperglycemic clamp to quantify beta cell function *in vivo*, we also established that beta cell function is not altered in PKO compared to control mice, as there were no statistical differences in GIR or hormone data between groups (**Figure 4**).

**Figure 3.**
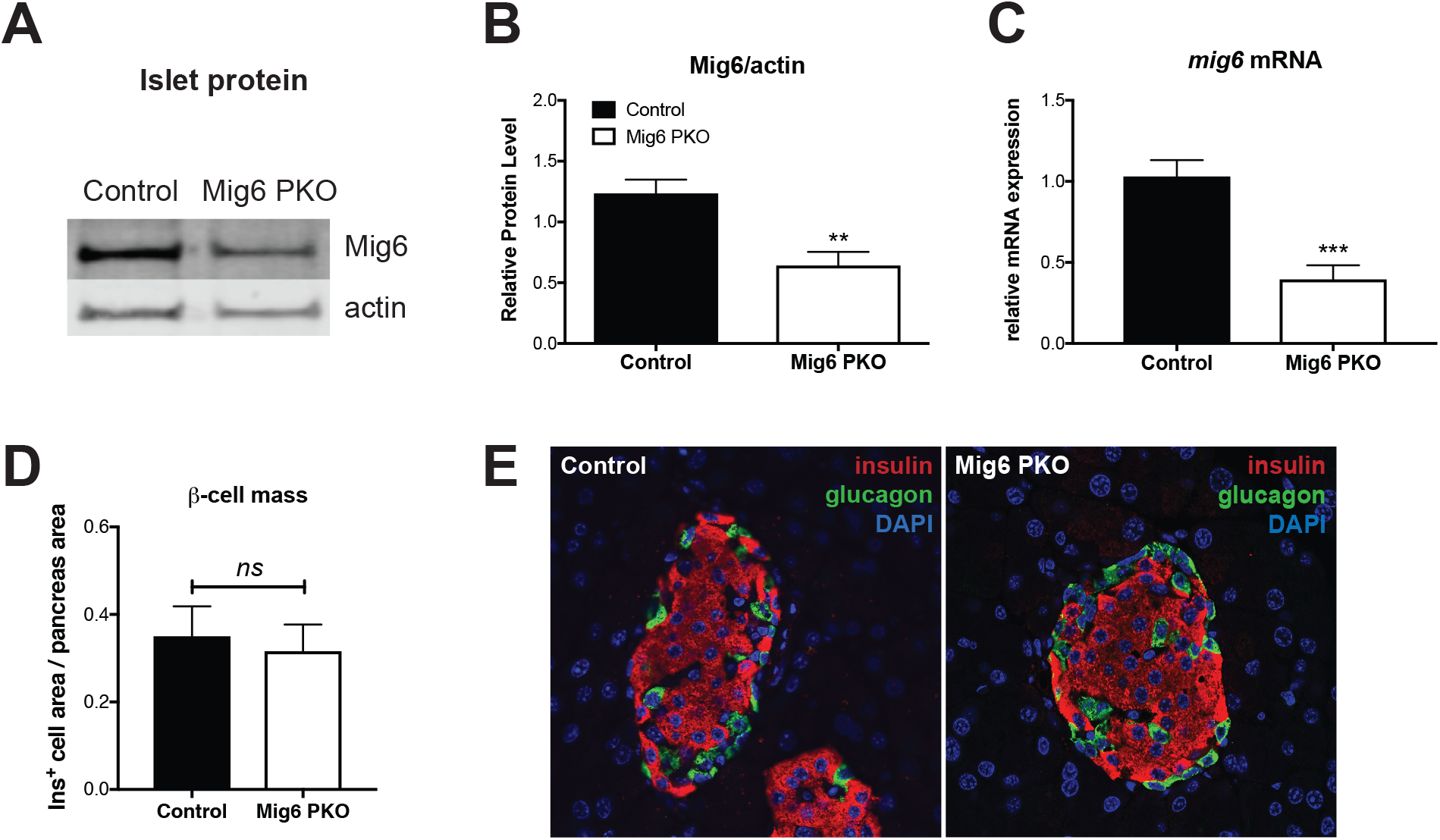
Pancreatic Mig6 knock-out (Mig6 PKO) mice have decreased Mig6 protein levels and mRNA expression. **(A)** Western blot representative image for graph **(B)** where islets were collected from control and Mig6 PKO mice and lysates collected for protein analysis. *n=6-7;* ***, *p<0.001 Student’s t-test*. Similarly, in **(C)** islets were isolated for cDNA preparation. *Mig6/Errfi1* levels were determined by qPCR and normalized to *Gapdh* relative to a chosen control sample. *n=3-5*; *, *p<0.05 Student’s t-test*. **(D)** Quantified islet area relative to total pancreatic area from saline-injected mice of both genotypes*. n=4-6*. **(E)** Images of immunofluorescent staining of insulin (red) and the glucagon (green), from an untreated control vs. Mig6 PKO mouse islet.

**Figure 4.**
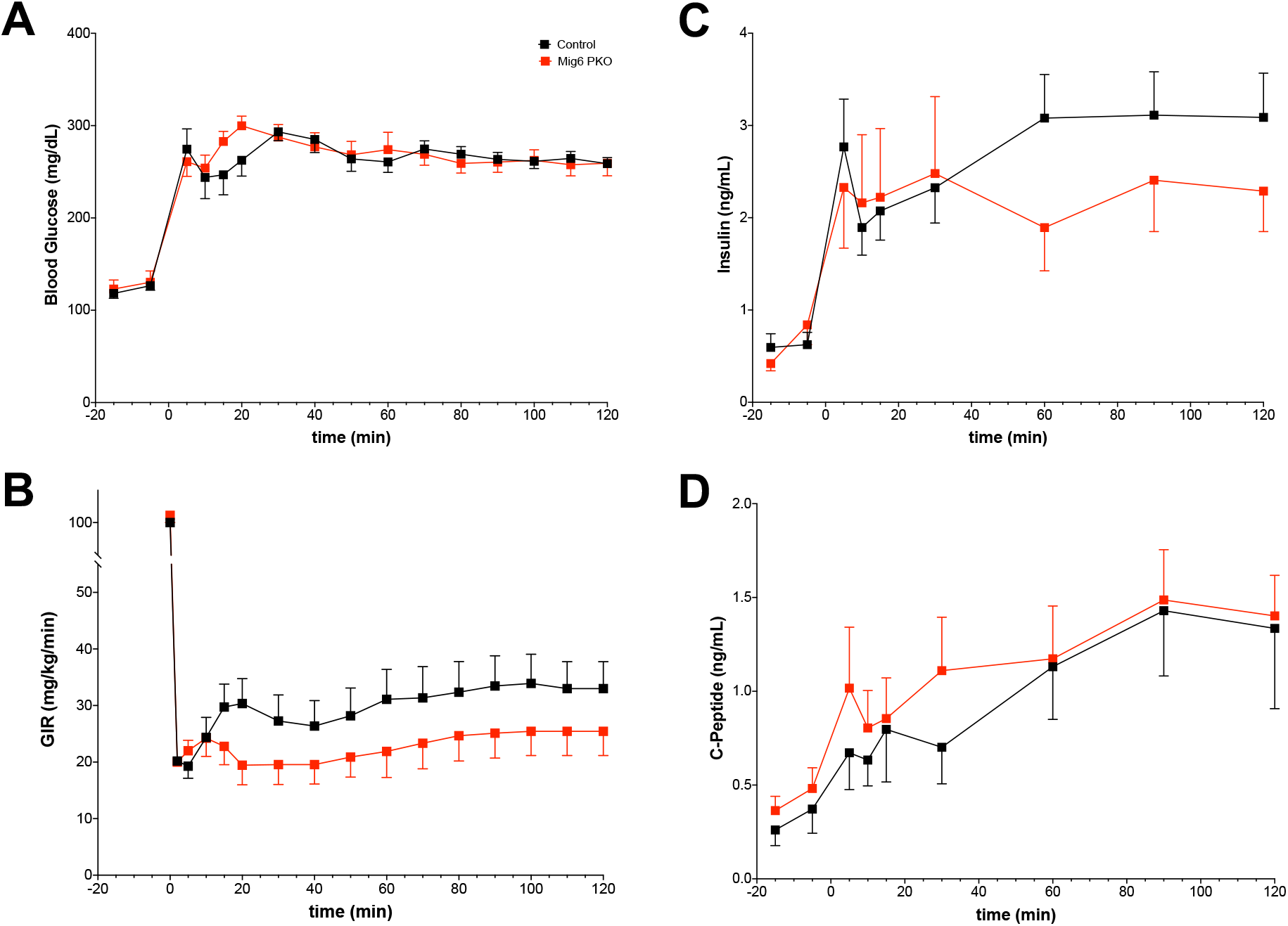
Insulin secretion *in vivo* in control and PKO mice. Insulin secretion was quantified in vivo using hyperglycemic clamps in control and PKO mice. **(A)** A primed followed by variable glucose infusion was used to clamp blood glucose at 270 mg/dl (15 mM). There were no differences between groups. **(B)** Glucose infusion rate (GIR) also did not differ between groups. Arterial blood was sampled throughout the procedure to depict basal, first-, and second-phase insulin secretion, which are derived from insulin **(C)** and C-peptide **(D)** concentrations. *n=7-8/group*.

### Mig6 PKO mice have lower fasting blood glucose and increased glucose tolerance after STZ treatment compared to control mice

Multiple low doses (MLD) of STZ causes hyperglycemia by damaging DNA in the beta cell and by inducing inflammatory mediators that infiltrate the islet, promoting beta cell dysfunction and death [35, 36]. MLD-STZ treatment is an experimental model that mimics selected features of the autoimmune-mediated beta cell destruction observed during the progression to human T1D. We subjected Mig6 PKO and control mice to MLD-STZ, then challenged them by performing glucose tolerance tests at 3-days and 20-days post STZ injection. Three-days post STZ treatment, whereas control littermates developed fasting hyperglycemia, Mig6 PKO mice maintained fasting blood glucose comparable to pretreatment blood glucose concentrations (**Figure 5A-B**). Mig6 PKO mice maintained this glycemic control to at least 20-days post STZ treatment. In addition, when given a glucose challenge, Mig6 PKO mice treated with STZ remained glucose tolerant compared to their littermate controls at 3-days post STZ treatment and this beneficial effect perpetuated 20-days post STZ (**Figure 5C-F)**.

**Figure 5.**
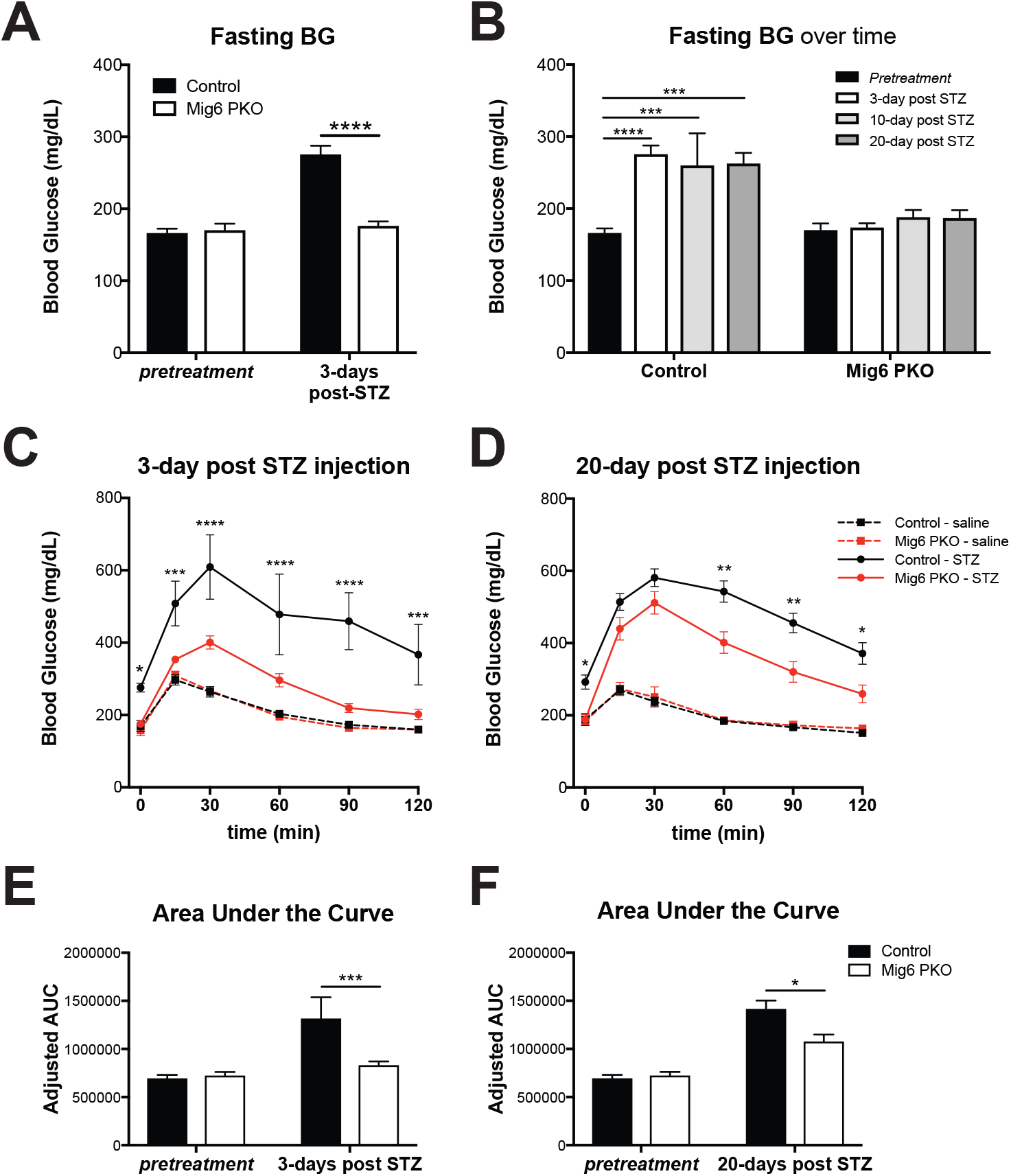
Mig6 PKO mice have a lower fasting blood glucose after STZ treatment, and have a slightly higher glucose tolerance than wild-type littermates. **(A)** 6-h fasting blood glucose (FBG) of both genotypes before and after treatment. *n = 3 (STZ CTRL) n=16-24 (other groups);* ****, *p<0.0001, 2-way ANOVA vs control*. **(B)** 6-hour FBG over time post-STZ treatment, showing that Mig6 PKO mice maintain FBG up to 20-days post STZ injection *n = 3 (STZ CTRL) n=16-24 (other groups);* ***, *p<0.001;* ****, *p<0.0001 in 2-way ANOVA vs control pretreatment*. **(C)** 10-week old Mig6 PKO and control mice received daily injections of 35 mg/kg body weight of streptozotocin (STZ) or volumetric equivalent of isotonic saline over 5 consecutive days. Intraperitoneal glucose tolerance test (GTT) of Mig6 PKO and control mice 3 days and **(D)** 20 days post-STZ or post-saline injection. **(E)** Calculated area under the GTT curve at 3-days and **(F)** 20-days post STZ injection.

### Mig6 PKO mice have preserved beta cell mass and enhance proliferation after STZ treatment

Prolonged fasting normoglycemia and glucose tolerance suggests Mig6 PKO mice are protected against STZ injury. We again performed IHC staining for insulin on 20-days post STZ sections and normalized to pancreas area. Beta cell crosssectional area was significantly higher in STZ-treated Mig6 PKO mice compared to STZ-treated control mice and was not different to saline-treated mice (**Figure 6A-B**). At 3- and 20-days post STZ treatment, we counted the total number of established islets, omitting clusters of less than 10 beta cells. The number of islets at 20-days post STZ treatment was doubled in PKO compared to control mice, and similar ratios existed at 3-days post STZ treatment but were not quite significant. Because there was no marked change in islet architecture, we suspected that beta cell proliferation contributed to the observed protection. We stained sections with Ki67, a marker of cells in the cell cycle. We counted the number of Ki67-positive beta cells in sections of control or Mig6 PKO mice. At 3 days recovery, Mig6 PKO mice had an increased proportion of Ki67-positive beta cells relative to total islet area (**Figure 6E**), but this proliferative effect ceased by 20 days of recovery (**Figure 6F**), suggesting that following STZ injury, Mig6 PKO mice have more proliferating beta cells in the early stages of recovery. Intuitively, this dip in proliferative response may contribute to the waning protection against hyperglycemia at 20 days recovery.

**Figure 6.**
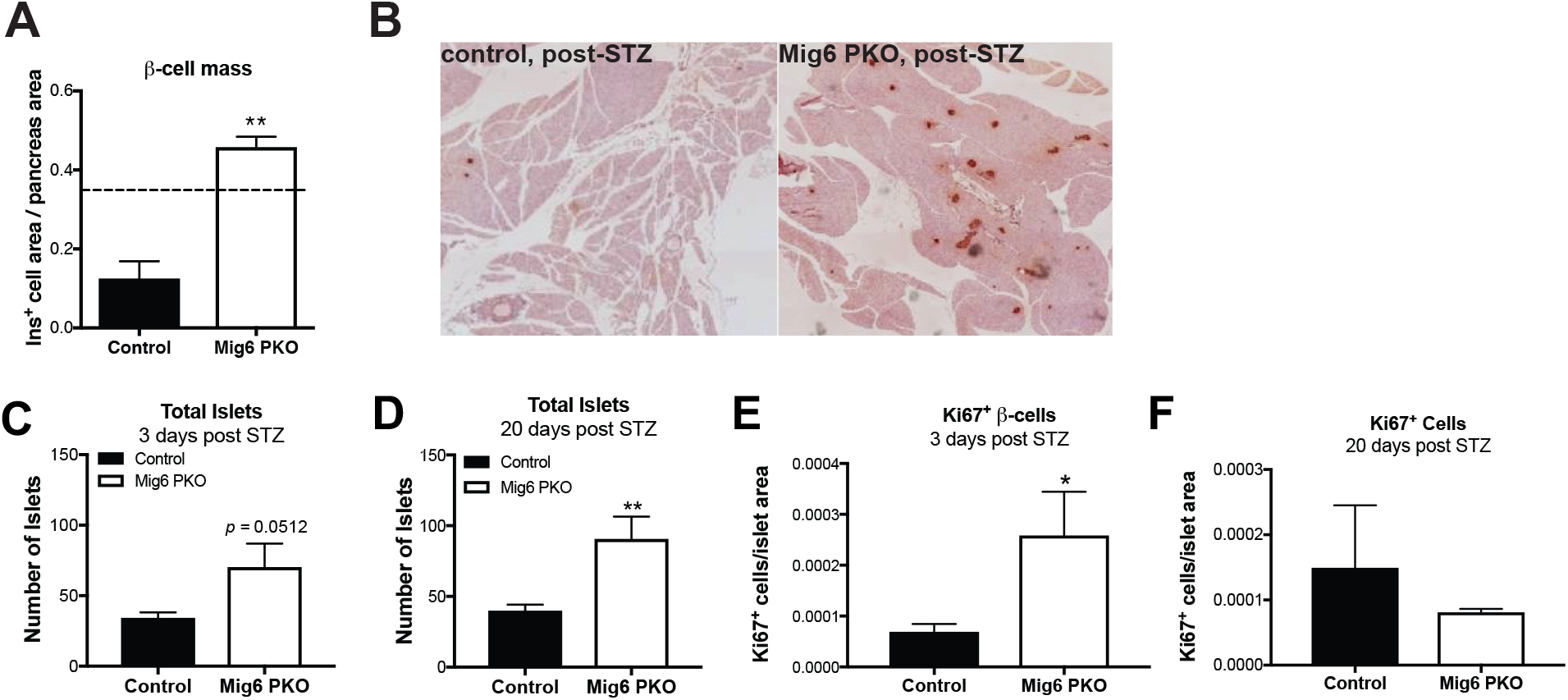
Mig6 PKO mice have preserved beta cell mass and increased proliferation. **(A)** Quantified islet area relative to total pancreatic area after STZ treatment (*dotted line represents saline-injected control mice*) calculated from **(B)** immunohistological staining for insulin in pancreatic sections of control and Mig6 PKO mice 20 days post-STZ treatment, *n = 4-6;* **, *p<0.01 vs control Student’s t-test*. **(C, D)** Immunofluorescent staining in pancreas sections of control and Mig6 PKO mice was employed to calculate total number of islets at 3- and 20-days post STZ, *n=3-5;* **, *p<0.01, Student’s t-test*. **(E, F)** Ki67-positive beta cells, detected by immunofluorescence, were counted and normalized to islet area at 3- and 20-days post STZ treatment. *n=3 each group;* *, *p<0.05 Student’s t-test, one-tailed*.

## DISCUSSION

EGFR signaling has been a focus of studies aiming to restore beta cell mass in patients with diabetes. Past studies have illustrated the potential benefits of cellular therapies: combinatorial administration of EGF and gastrin [8] rescues hyperglycemia, and constitutively active EGFR increases beta cell mass in mice [14]. In this study, we examined how pro-inflammatory cytokines, widely accepted to participate in the immune response during T1D, restrain EGFR signaling. Cytokines induce iNOS expression and NO production, and we demonstrated that cytokine-induced NO is necessary for Mig6 expression. We also present evidence that EGFR signaling is dampened by NO, from cytokines or an NO donor itself. We suggest that the induction of Mig6 with cytokines, which in turn, inactivates EGFR signaling, is deleterious to the beta cell’s efforts to mount a recovery response.

To advance our work *in vivo* and build upon our previous mouse model of Mig6 haploinsufficiency, we generated a pancreas-specific knock out of Mig6 and observed a comparable, but more durable phenotype to the Mig6 haploinsufficient mouse model. Under normal conditions, the metabolism of control and PKO mice was indistinguishable and major differences in islet architecture or beta cell mass were absent. In addition, beta cell function *in vivo* was unaltered in PKO mice compared to their control littermates. When challenged with a very low dose STZ model, designed to simulate the early events of beta cell dysfunction and destruction in T1D, control animals develop fasting hyperglycemia and glucose intolerance. Strikingly, Mig6 PKO mice have normal fasting blood glucose which persists at 3-, 10- and 20-days post STZ treatment; Mig6 PKO mice also present improved glucose tolerance compared to their littermate controls after STZ treatment. Most notably, Mig6 PKO mice have increased beta cell mass 20 days post STZ compared to their wildtype littermate control mice. These data suggest that Mig6 may aid in the progression to diabetes after initial injury to the beta cells.

During our histological studies, we identified that Mig6 PKO mice had increased numbers of Ki67-positive, proliferating beta cells. Our previous report in the Mig6 haploinsufficient mouse model concluded that there was no difference in STZ-stimulated beta cells that have entered into the M phase of the cell cycle. Here, we report a more encompassing marker of proliferation, Ki67, which identifies all cells in any stage of the cell cycle. Because it is suggested that duct-associated islets and single extra-islet beta cells indicate ongoing duct-to-islet neogenesis [37], we also noted the number of islets (including singlets) that were associated with a duct. Numbers of duct-associated islets were not different between the control and Mig6 PKO mice (data not shown), indicating an intrinsic change in cell fate. We defined the mechanism of beta cell mass preservation as exploitation of beta cell proliferation, which occurred in the early days following beta cell destruction with STZ. The involvement and contribution of other means of beta cell restoration, such as beta cell transdifferentiation and neogenesis, which in other transgenic models have been shown to potentially increase the reservoir of new beta cells [38, 39], have yet to be addressed in the context of Mig6 knock down.

It might seem surprising that deletion of an anti-proliferative factor such as Mig6 did not promote beta cell mass expansion alone. However, given that Mig6 is a feedback inhibitor of EGFR, beta cells would still require a signal (e.g., EGFR ligands) to drive replication. We view this requirement for endogenous (or even addition of exogenous) mitogenic factors as beneficial from a safety standpoint (i.e., pancreatic Mig6 deletion alone is not tumorigenic). Indeed, we have not observed pancreatic tumors in any of the mice studied to date, albeit we have not conducted prolonged studies. Nevertheless, coupling Mig6 inhibition or deletion with EGF (or other EGFR ligands) could have greater translational potential than either strategy alone and increase the robustness of a protective effect following beta cell attack.

In summary, our data demonstrated that NO is detrimental to EGFR activation and phosphorylation, and is required for cytokine-mediated Mig6 expression. Yet NO alone does not induce Mig6. As in our previous Mig6 haploinsufficient mouse model, when suppression of Mig6 in the pancreas protects mice from chemically-induced hyperglycemia by preserving beta cell mass. This study highlights the pitfalls of treatments solely focused on the immunological response, as pathological stimuli can detrimentally impact mitogenic signaling and therefore EGFR repair mechanisms via Mig6 feedback inhibition. Mig6 presents a promising, adjunct cellular therapeutic alternative that, when used to target beta cell recovery programs, may prevent or reverse the progression to diabetes.

## ACKNOWLEDGEMENTS

We would like to thank K. Benninger and J. Nelson (Indiana University Center for Diabetes and Metabolic Diseases) for assistance with islet isolation and imaging. Thanks to Dr. Andrew Lutkewitte, PhD (Washington University, St Louis, USA) for thoughtful discussions and assistance with husbandry.

---

**Supplementary Data Table 1.** List of antibodies for immunostaining

| **Name** | **Vendor, catalog number** | **Dilution** |
| --- | --- | --- |
| Anti-insulin | Santa-cruz, #H-86 | 1:250 |
|  | Invitrogen, #180067 | 1:250 |
| Anti-glucagon | Abcam, #ab92517 | 1:500 |
| Anti-Ki67 | Abcam, #ab15580 | 1:900 |

