## Supplemental Table 1 for "Pancreatic Deletion of Mitogen-inducible Gene 6 Promotes Beta Cell Proliferation Following Destruction"

Supplementary Data Table 1. List of antibodies for immunostaining

| **Name** | **Vendor, catalog number** | **Dilution** |
| --- | --- | --- |
| Anti-insulin | Santa-cruz, #H-86  Invitrogen, #180067 | 1:250  1:250 |
| Anti-glucagon | Abcam, #ab92517 | 1:500 |
| Anti-Ki67 | Abcam, #ab15580 | 1:900 |
